## Supplemental Figures for "A draft genome assembly of the agricultural pest *Leucoptera coffeella* and analysis of its dsRNA processing machinery is a key step towards RNAi-based biopesticides in Lepidoptera"

**Supplemental Materials**

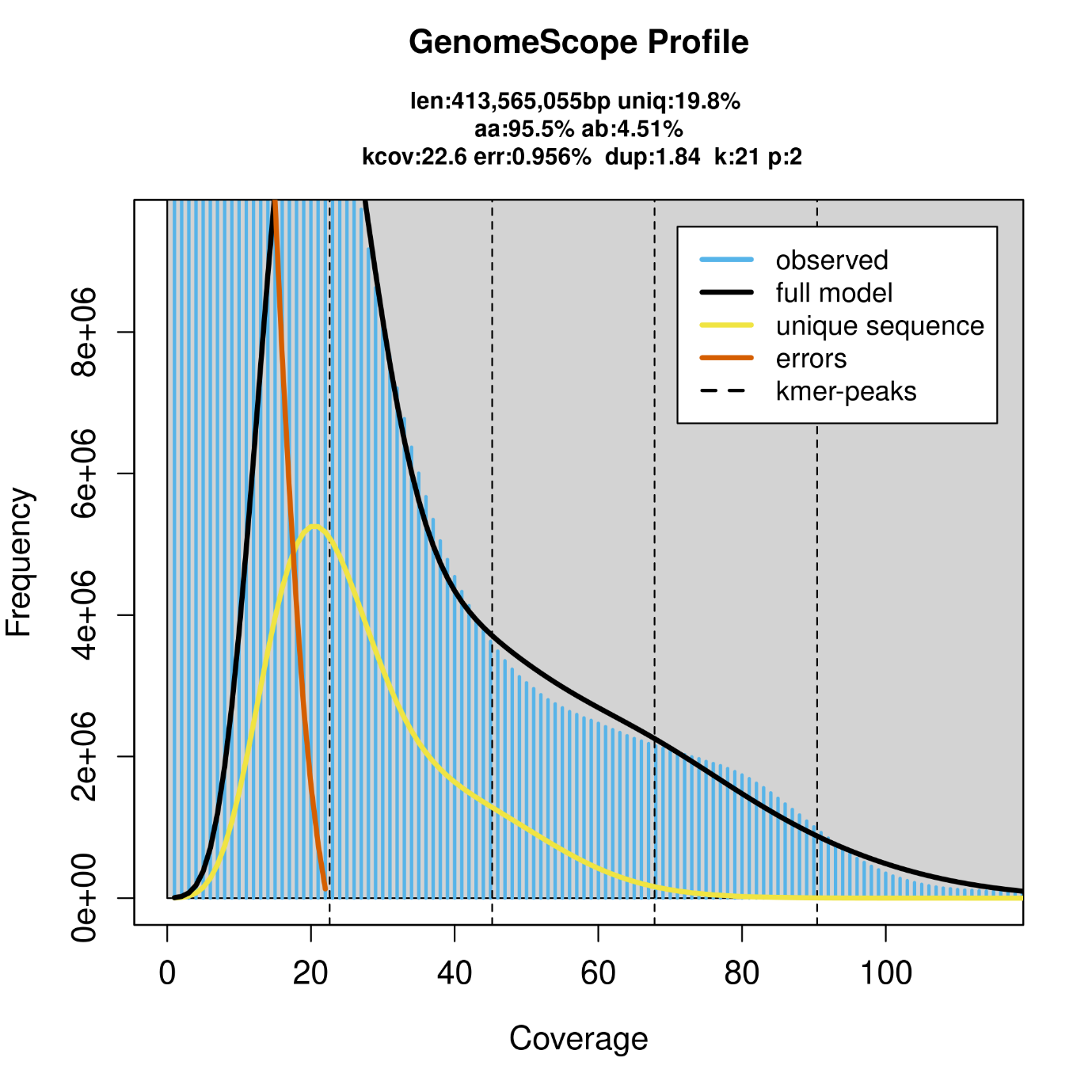

**Figure S1.** Coverage plot produced by jellyfish and GenomeScope 2.0 analysis of our raw reads. The pooled sample generated a large amount of low coverage kmers due to the genetic heterogeneity present in our sample.

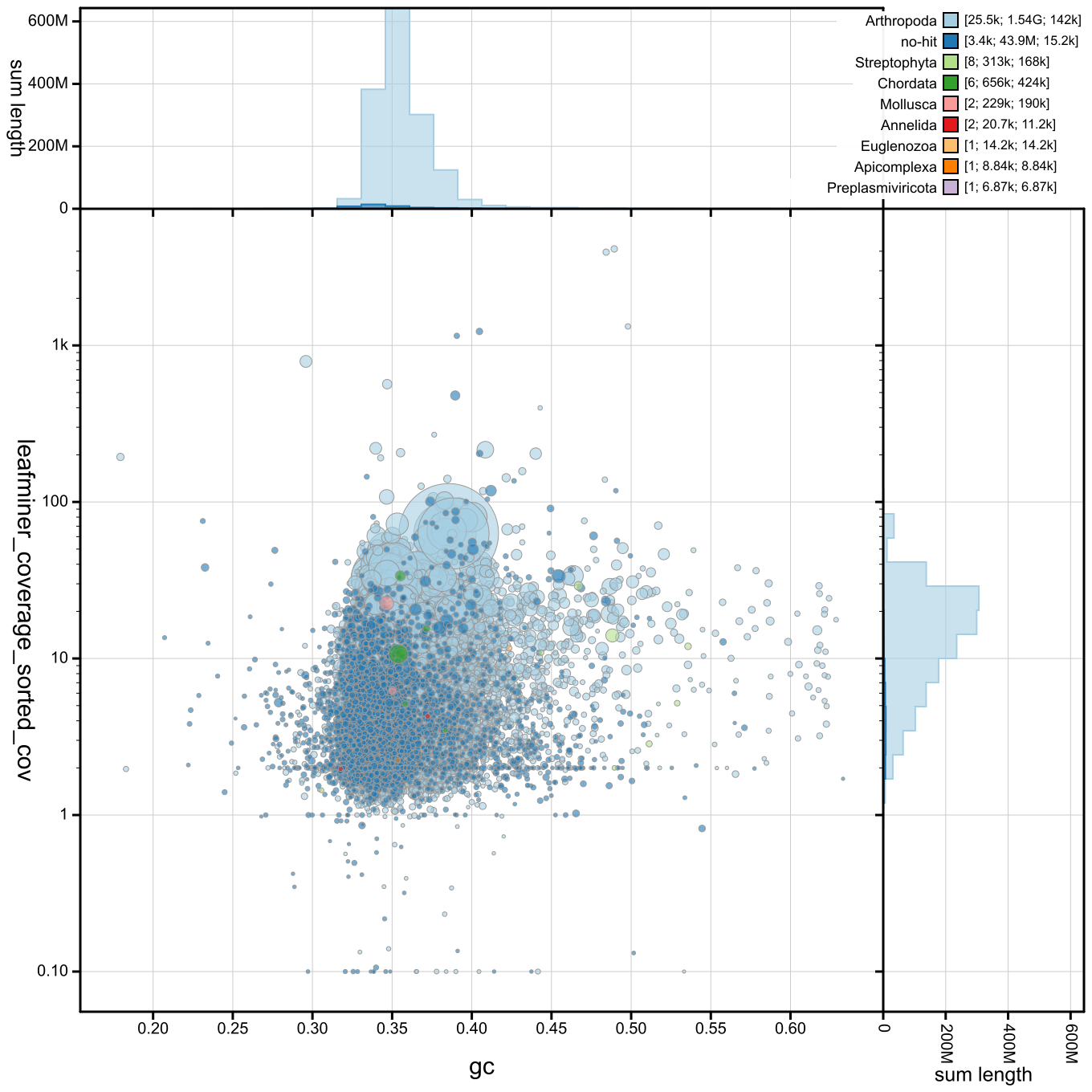

**Figure S2.** Blobplot of preliminary metaMDBG assembly. The vast majority of contigs were identified as Arthropoda, although a variety of low coverage contigs were identified as contamination.

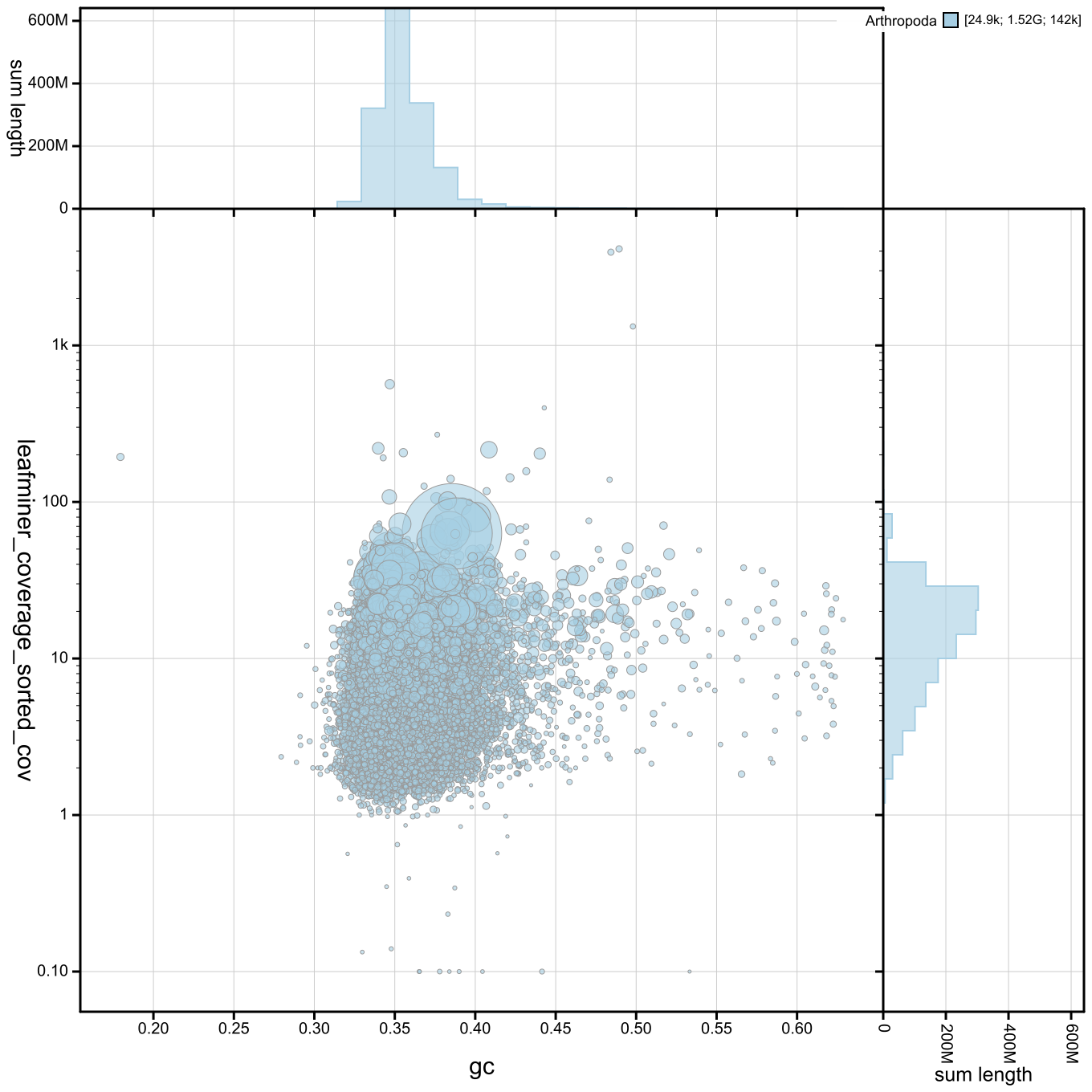

**Figure S3.** Blobplot of metaMDBG assembly after filtering to include only Lepidoptera contigs using blobtoolkit filter function. All remaining sequences belong to phylum Arthropoda.

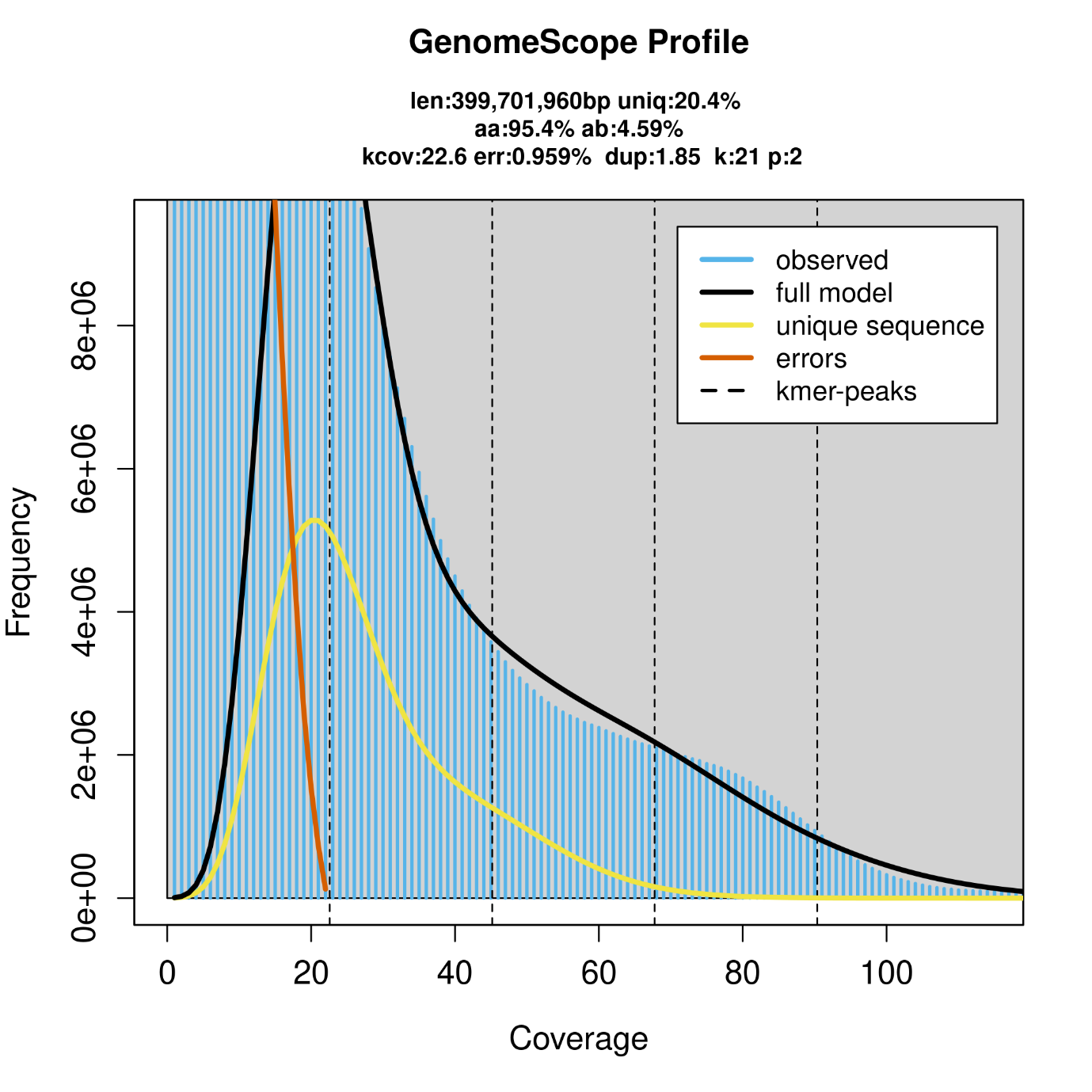

**Figure S4.** Coverage plot produced by jellyfish and GenomeScope 2.0 analysis on the filtered reads (Lepidoptera only). The large amount of low coverage kmers are still present, indicating that they are due to genetic variation within our pooled sample rather than the presence of contaminant reads.

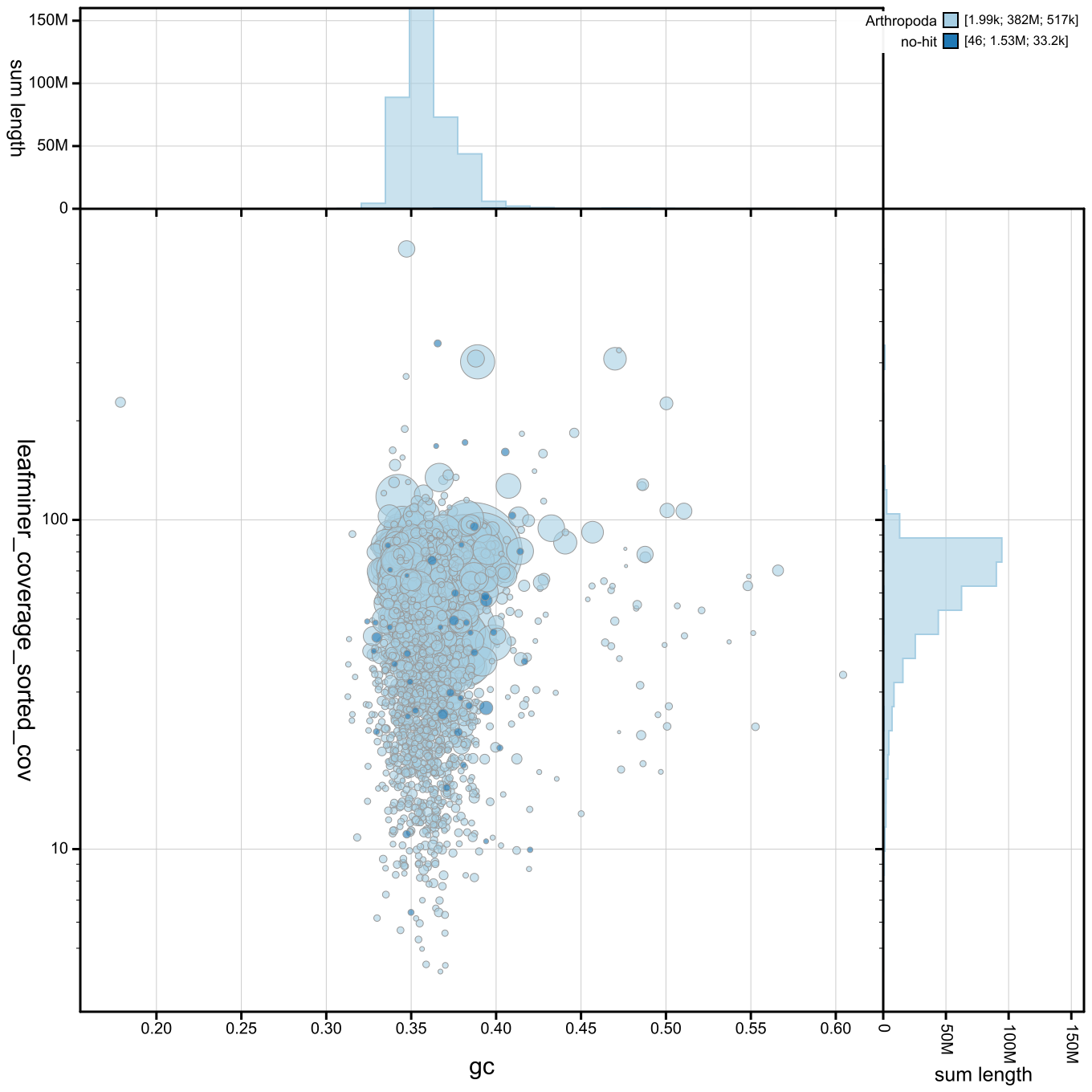

**Figure S5.** Blobplot of hifiasm assembly produced using only the reads that mapped to Lepidoptera contigs in our preliminary metaMDBG assembly.

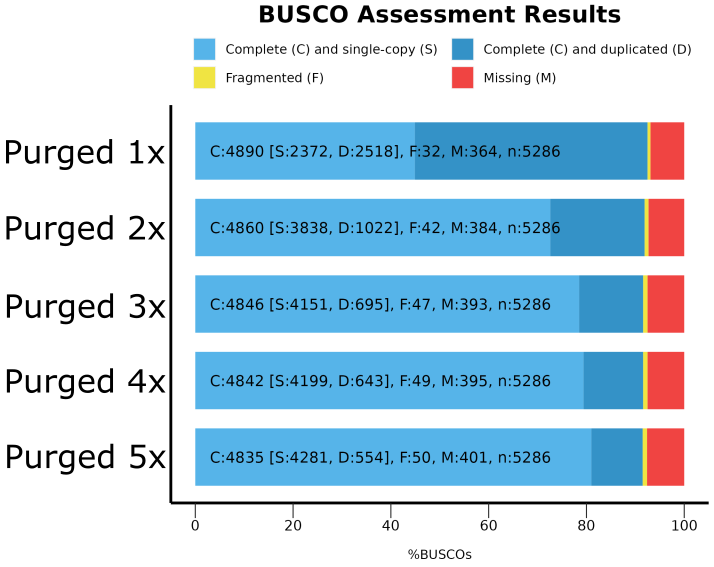

**Figure S6.** BUSCO plot showing intermediate assemblies after iterative rounds of the purge_dups algorithm. After 5 rounds of purging, the duplicated BUSCO count was reduced from 2518 to 554 while only modestly increasing the number of fragmented (32 to 50) and missing (364 to 401) BUSCO genes.

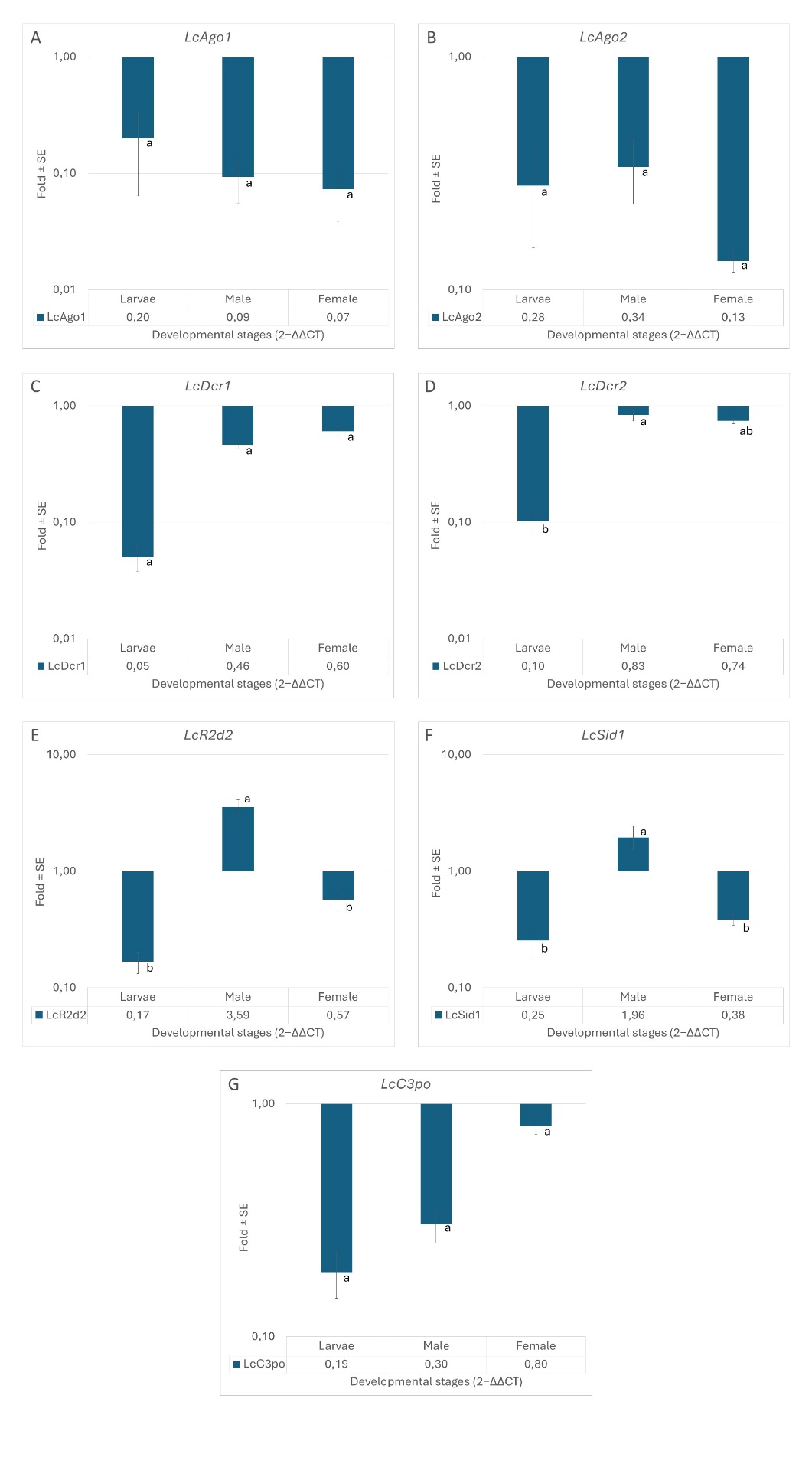

**Figure S7.** Expression of *L. coffeella* RNAi genes as measured via qPCR.

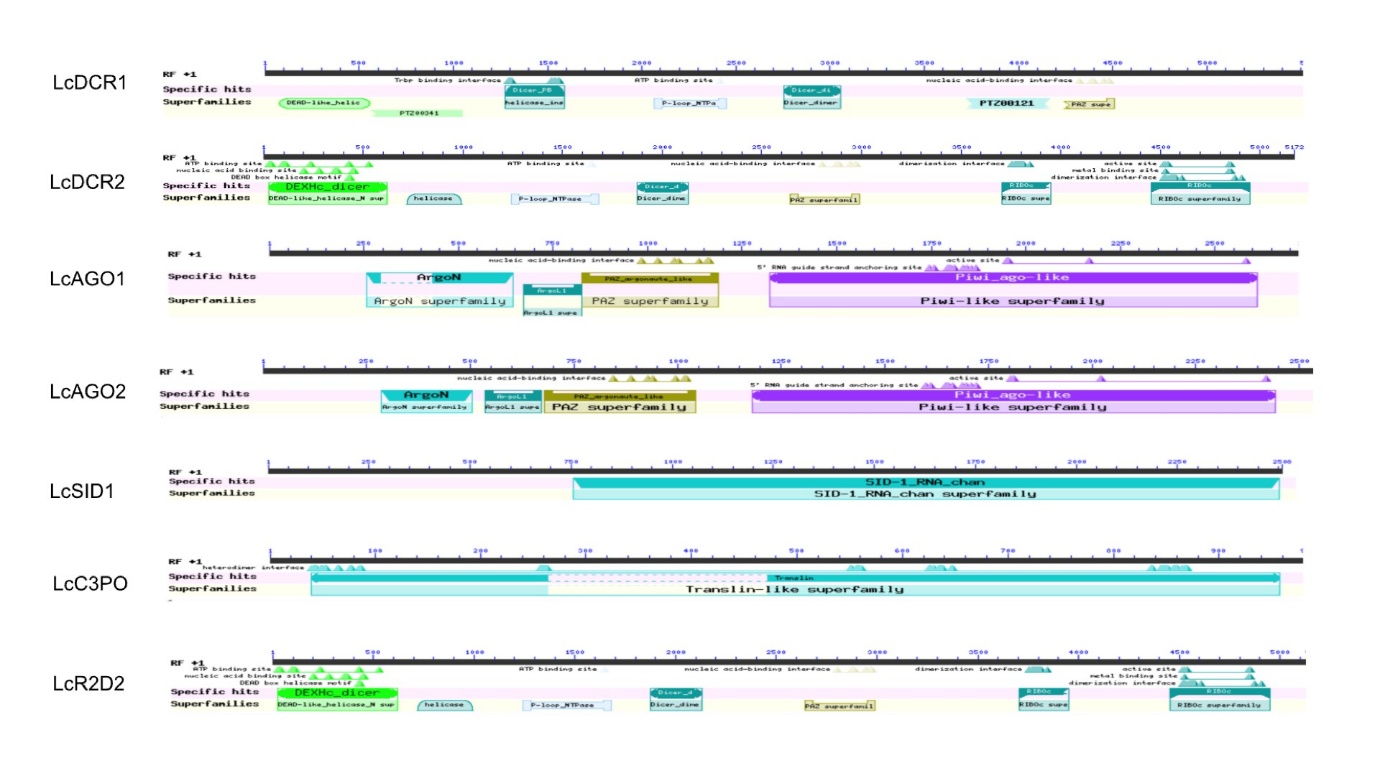

**Figure S8.** Gene models of *L. coffeella* RNAi genes identified within our genome.

**Table S1:** qPCR primers sequences

**Table S2.** Detailed assembly statistics generated by Inspector before and after polishing

**Table S3.** Detailed RNA-seq mapping statistics

**Table S4.** Repetitive element content as measured by RepeatMasker
